# Co-infection with two iflaviruses (deformed wing virus and sacbrood virus) affects viral and immune dynamics and synergistically increases honey bee mortality

**DOI:** 10.1101/2024.04.23.590860

**Authors:** Alice Mélusine Durand, Eric Dubois, Anne Bonjour-Dalmon

## Abstract

The honey bee microbiome includes a wide variety of viruses. While most of them usually remain commensal, some can become pathogenic in specific contexts. Of these, one is that of deformed wing virus (DWV) and another, sacbrood virus (SBV). Although co-infection is the norm rather than the exception, most of the time these viruses have been studied independently. When investigated as co-infections, past studies have focused on their effects on the honey bee brood. In this study, we co-inoculated adult honey bees at emergence with DWV by injection and SBV orally (acting as the viral transmission by *Varroa destructor* and by trophallaxis or food, respectively), either simultaneously or sequentially. Using optical counters, we were able to track the survival and behaviour of these honey bees within colonies. Through regular in-hive sampling, we monitored the evolution of their viral loads as well as the expression of eight immune genes involved in honey bee anti-viral immunity. Here, we show that co-inoculations of DWV and SBV synergistically increase the virulence of DWV and conditionally promote the replication of both viruses. Finally, our results show that immune responses in adult honey bees depend on DWV genotypes and whether replication originates from a superinfecting virus or a virus already present in bees.

**Author Summary:** Honey bees are highly social pollinators that live in crowded colonies. Their population density and the high frequency of interactions between individuals favours disease transmission and makes colonies susceptible to pathogen outbreaks. Many viruses commonly infect honey bees, however, they are often studied as single infections. As an effort to better understand interactions between honey bees and multiple viral populations, we co-inoculated young bees with two common honey bee viruses (deformed wing virus and sacbrood virus), released them in colonies and monitored their health and behaviour. Our findings show evidence of synergies between both viruses, as we show that a virus seemingly harmless for adult bees (sacbrood virus) may actually increase the virulence of another virus (deformed wing virus). These results highlight the importance of monitoring and studying multiple pathogens at once for a better understanding of the threat they represent to colony health and survival.

## Introduction

Many different viruses are capable of infecting honey bees [1]. Among them, deformed wing virus (*iflavirus aladeformis*, DWV) and sacbrood virus (*iflavirus sacbroodi*, SBV) have become serious threats to honey bees all over the world [2–4].

Before the spread of *Varroa destructor* (an ecto-parasitic mite of honey bees), DWV was mainly transmitted horizontally between bees through feeding or trophallaxis [5,6], vertically from queens to eggs [7,8] and during mating [9]. Since the varroa mite switched from *Apis cerana* (the Asian honey bee) to *Apis mellifera* (the western honey bee) [10], it has gradually been introduced to most parts of the world [11–15]. Varroa reproduces by infesting capped brood and feeding on pupae haemolymph and fat body tissues [16], but it also feeds on adult bees [17]. When feeding, varroa can transmit various viruses to honey bees [18–20]. This additional route of infection has greatly increased the pathogenicity of DWV [19,21], especially when combined with the ability of some DWV strains (DWV-B) to replicate within the mite [22,23]. This combination first led to a decrease in DWV genetic diversity [24], then a new diversification of DWV strains through competition [25,26] and recombination [27,28]. DWV-B has been found to be more virulent than strains that do not replicate within the mite (DWV-A) [19,29,30] and more efficient in its replication [31], leading to gradual replacement of DWV-A by DWV-B [32].

Conversely, SBV dynamics and pathogenicity have not been directly affected by this new potential route of transmission by varroa [20]. However, the mite may still indirectly impact SBV dynamics as SBV has been found to be more prevalent and responsible for higher loads in bees from colonies infested by the mite [33], with a positive correlation between SBV loads in mites and adult bees [34]. In occidental countries, SBV infections are almost ubiquitous. Although this situation does not appear to be a critical threat to *Apis mellifera* colonies [35], the potential danger of this virus should not be overlooked. While most SBV strains represent a major threat for *Apis cerana* colonies [36,37], the pathogenicity of SBV can vary in *Apis mellifera* depending on SBV strains [38], suggesting that a small genomic variation can result in differential pathogenicity. Given the high mutation rate of RNA viruses [39], SBV should therefore be carefully monitored as it has the potential of becoming a serious threat to *Apis mellifera* as well.

While both viruses are increasingly studied as single infections, only a handful of studies have investigated co-infections with SBV and DWV [40,41], and all of them have focused on effects and dynamics in honey bee larvae and pupae only. While symptoms related to these viruses are indeed associated with infections at early stages of development, adult bees still often carry high viral loads [42–44]. The pathogenicity of different DWV strains has been studied in adult bees by some authors [19,29], as well as potential sublethal effects of these infections [45–47]. Such sublethal effects may lead to a decrease in the foraging potential of the colony, which in turn may increase overwintering mortality or reduce honey production. To our knowledge, interactions between the two viruses and the consequences of one viral infection on the dynamics of the other virus have never been studied in adult bees living in hives.

In this study, we tackled the question of whether DWV and SBV interact in adult honey bees living in their colonies and investigated potential sublethal effects of both viruses, either alone or in co-inoculation. In an effort to reflect plausible scenarios of infection and bring our study closer to natural conditions, we studied both simultaneous and sequential co-inoculation of DWV and SBV into bees living in hive colonies. To better understand the dynamics of both viruses *in situ* and their consequences on honey bee health and daily activities, we analysed the behaviour and survival of co-inoculated bees and quantified six major honey bee viruses. Additionally, virus co-inoculations may impact the honey bee immune system differently than single inoculation, thus triggering immune-mediated viral interactions (reviewed in [48]). As such, we also quantified the transcription of eight immune genesinvolved in various honey bees’ anti-viral immunity pathways, namely the RNAi pathway (*argonaute-2* and *dicer*), the Imd pathway (*relish*), the humoral pathway (the anti-microbial peptides (AMP) *defensin-1* and *defensin-2*), the melanisation pathway (*prophenoloxidase*), the Jak-STAT pathway (*vago*) as well as *vitellogenin*, which expression is involved in bee longevity and immune potency.

## Material and methods

### Virus preparation and inoculation

One strain of each virus (SBV, DWV-A and DWV-B) was used for the experimental inoculation of honey bees. The strains were selected from archived honey bee samples (supernatants of symptomatic bees in 10 mM phosphate buffer pH 7) at the ANSES laboratory of Sophia Antipolis on the basis of their previous viral quantification [49]. The samples were selected only if the quantification of the virus of interest was higher than the quantification for ABPV, CBPV, BQCV and DWV or SBV (at least 10^7^ times higher for DWV samples; at least 10^10^ for SBV samples). The genotype of the selected DWV-A and DWV-B samples had previously been analysed by next-generation sequencing [49]. The DWV-A sample (MD-GhidighiciChisinau23_2017_H156) originated from bees collected in Moldova in 2017 and was found genetically pure. The DWV-B sample (ES-Arbeiza22_2017_H62) originated from bees collected in Spain in 2017 and contained pure DWV-B as well as recombinants containing the IRES region from DWV-A. The SBV strain originated from bees collected at Sophia-Antipolis, France, in 2017. All the strains were multiplied through injection into white-eyed pupae. Seven days post-injection, the pupae were individually crushed in PB (1 mL/pupa) and centrifuged for 10 min at 8,000 x *g* at 4°C. The supernatant was then transferred to a new tube, and centrifuged again. Supernatant from this second centrifugation was aliquoted and analysed for the six major honey bee viruses by quantitative reverse transcription PCR (RT-qPCR) after RNA extraction [50]. Aliquots were filtered through a 22 nm filter before being used as inoculums for the experiments.

DWV was inoculated through injection (Nanoject II - Drummond Scientific, Broomall, PA, USA) between the third and fourth tergites of emerging bees. DWV inoculums were diluted in PBS so as to inject 1•10^6^ genome copies in 46 nL. Honey bees from the same experimental condition were collectively anaesthetized with CO_2_ and kept on ice until they were all injected. In contrast, SBV was inoculated orally. SBV inoculums were adjusted in a 30% (w/v) sucrose solution so as to feed 1•10^8^ genome copies in 5 µL. Honey bees were starved for 2 h before being handled individually and fed using a micropipette. Sucrose feeders were placed back in the bee cages between 30 min and 2 h after inoculation. Honey bees that endured both methods on the same day were always injected first (DWV or PBS), then fed (SBV or sucrose) to avoid regurgitation during anaesthesia or injection. Inoculums were diluted on day 0 for each week of experimentation and kept on ice in a refrigerator (4°C) until they were discarded after day 2 inoculations (see Section 2.2.).

### Experimental design

Our experiments were conducted between April and June 2022 and in April 2023. During March 2022 and 2023, pools of 40 frame-bees from our experimental apiary colonies were tested for the six major honey bee viruses (ABPV, BQCV, CBPV, DWV-A, DWV-B and SBV) by RT-qPCR. The three colonies showing the lowest viral loads were transferred into small hives connected to an optical bee counter [51] (henceforth: ‘host colonies’). For each year of experimentation, three colonies with low viral loads were also selected as a source of emerging bees (henceforth: ‘donor colonies’). In these experiments, interaction between DWV-A and SBV and interaction between DWV-B and SBV were tested. Both interactions were tested over two years in 3 colonies each (S2 Table). Due to the large number of bees required, each experiment (one interaction in one host colony) was divided up and performed over the course of two weeks, which were either consecutive or one week apart.

To assess potential effects of the chronology of infections on virus interactions, inoculations were performed on day 0 and/or day 2 of each week of experimentation (Fig 1). The two viruses were either inoculated simultaneously on day 0 or 2, or sequentially on day 0 and 2. Sets of honey bees were also inoculated with a single virus (SBV, DWV-A or DWV-B) on day 0 or 0. To avoid potential methodological biases, multiple control treatments were included. Some sets of honey bees were fed with a sucrose solution (30% w/v) on day 0 or 2. Other sets of honey bees were fed with a sucrose solution (30% w/v) and injected with a PBS solution either simultaneously on day 0 or 2 or sequentially on day 0 and 2. One set of honey bees was not treated at all. In addition to bees injected with PBS and fed with sucrose, and so as to control for potential crossed effects of an orally inoculated virus and the injection method, the experiments conducted in 2023 tested sets of honey bees that were both inoculated with SBV and injected with PBS either simultaneously on day 0 or 2 or sequentially on day 0 and 2 (Fig 1).

**Figure 1.**
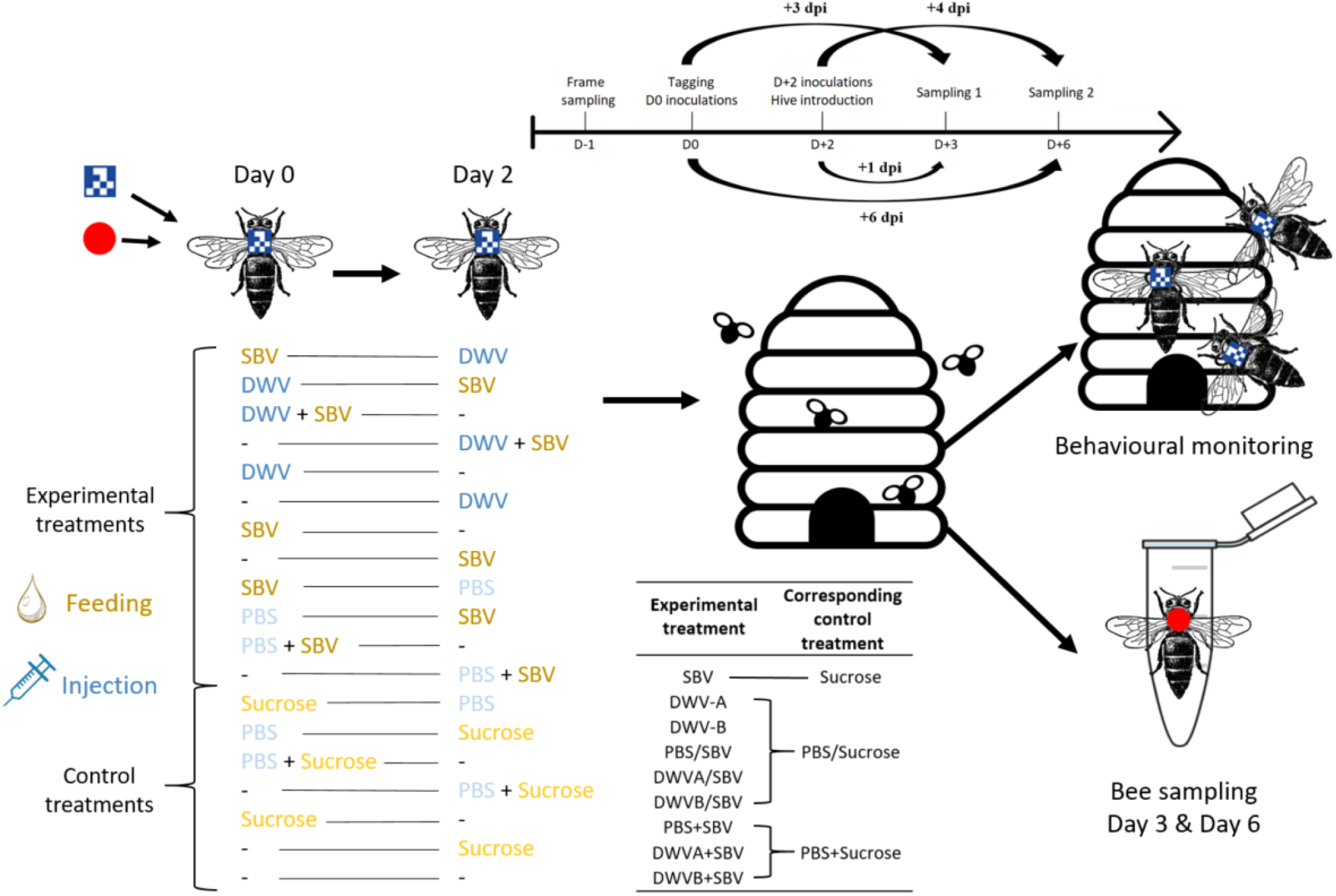
Schematic representation of the experimental protocol detailing bee treatments, their controls, and the chronological procedure. The “/” symbol represents sequential treatments on two different days while a “+” symbol represents simultaneous treatments on the same day.

The day before experimentation started, donor colony frames — from which adult bees had been removed and on which emerging brood was observed — were sampled and kept overnight in a dark incubator at 34°C and saturated humidity. On day 0, all the emerging bees were pooled together. A specific treatment was assigned to each cage and each paint colour. For each treatment, a set of honey bees was painted on the thorax using a POSCA^®^, while another set was marked with a 3-mm data-matrix QR code glued (Sader) on their thorax (S1 Fig). Any bee showing altered motility after marking was discarded. Day 0 inoculations were performed once all the bees had been marked. After inoculations, honey bees were kept in a dark incubator at 31°C and saturated humidity until day 2. After day 2 inoculations, all the bees were released into a host colony. On days 3 and 6, nine painted bees from each treatment were taken from the colony and flash-frozen on dry ice, then gathered into three pools of three honey bees and kept at −80°C until analysis. Since only one host colony per interaction was tested for treatments including both SBV inoculation and PBS injection, eighteen painted bees were sampled and gathered into six pools of three honey bees for each experiment. All the treatments and experimental procedure are summarised in Fig 1.

### Viral quantification and immune gene expression

Each pool of three honey bees was crushed with a 0.8 cm tungsten bead in 900µL of Trizol (Quiazol, Qiagen) using a Tissue Lyser II (Qiagen) and centrifuged at 11,000 x *g* for 30 s. A volume of 500 µL of supernatant was then transferred to a new tube. RNA was extracted according to the manufacturer’s recommendations (RNeasy Universal Mini Kit, Qiagen). A supplementary step was added between the first and second washing buffer to eliminate residual DNA (DNase PureLink, Invitrogen) according to the manufacturer’s instructions. The quantity of extracted RNA was estimated using a spectrophotometer (Nanodrop 2000, Thermo Fisher Scientific). Each sample was adjusted in H_2_O to 500 ng/µL of RNA. Reverse transcriptions were performed from 1 µg RNA according to the manufacturer’s protocol (High-capacity RNA to cDNA kit, Applied Biosystems), using random priming. Each plate was heated to 37°C for 60 min and 95°C for 5 min in a thermocycler (Eppendorf Mastercycler, Nexus SX5). Each well was then diluted ten times so as to obtain an estimated cDNA concentration of 50 ng/µL.

Six major honey bee viruses were quantified for each sample: DWV-A, DWV-B, SBV, ABPV, CBPV and BQCV. qPCR assays were performed in duplicate in a StepOnePlus real-time PCR system (Applied Biosystems, Life Technologies). In 96-well plates, 3 µL of diluted cDNA diluted was added to 1 µL of each primer (S1 Table) with 10 µM and 5 µL of SYBR Green Master Mix (Applied Biosystems, Life Technologies), using the following thermal program: 10 min at 95°C followed by 40 cycles comprised of 15 sec at 95°C and 1 min at 60°C; a final cycle comprised of 15 sec at 95°C, 1 min at 60°C and 15 sec at 95°C was used for melt curve determination. The quantification cycles (Cqs) were retrieved and referred to a range of seven dilutions of synthetic cDNA fragments of known quantity (Eurofins, France) to deduce target quantity in tested samples. Two H_2_O-based negative controls were systematically added to each plate: one control from the reverse-transcription step and one qPCR control.

The transcription of eight immune genes involved in various immune pathways was also quantified. To quantify their mRNA, a similar qPCR procedure was conducted on a Biorad CFX-384 in 384-well plates. The list of primers used in this quantification can be found in S1 Table. One H_2_O-based qPCR control was systematically added to each plate and for each target. PCR Cqs were retrieved and compared against two reference genes (RpL32 and RpS5). Additionally, a standard sample common to all the plates was added to account for any variability between runs and used as a reference to normalize variation. For each sample, all gene transcriptions were analysed on the same plate.

### Behavioural monitoring

In these experiments, a total of 7,241 bees were tagged with a data-matrix QR code (Table 2). Host colonies were connected to an optical counter [51] composed of a camera that recorded a modified hive entrance from above. The modified entrance was composed of eight tunnels that did not allow bees to walk over one another. As the bees were marked, the real-time monitoring software was able to identify individual honey bees, record the precise time and date of detection and infer the bee’s direction (going into or out of the hive; S1 Fig). Depending on the optical counter used, between 2% and 8% of recorded images did not allow identification. From this raw data, we were able to determine the proportion of bees that were detected, the pool of available and effective foragers, the age of each bee at its onset of foraging, survival after its first foraging trip, and the duration of each trip. A bee was considered an available forager if she completed at least one successful trip outside the hive and back inside. Contrastingly, a bee was considered an effective forager if she completed at least one trip outside the hive lasting more than 10 min. The last record of a given bee was considered as its time of death.

### Statistical Analysis

All statistical analyses were performed in R v4.2.2 (RStudio build 576). Data from each treatment group in each dataset were tested for normality (Shapiro-Wilk test, package *‘stats’*). As no dataset followed a normal distribution, non-parametric tests were used. As recommended by the OECD, the proportion of bees that died before reaching their onset of foraging was expressed as per cent of their respective controls [52]. Mortality proportions were compared through a pairwise nominal independence test with Benjamini-Hochberg adjustment acting as a post-hoc chi-squared test (package *‘rcompanion’*). Uncorrected mortality proportions were used to compare treatment groups to their respective controls through chi-squared tests (package *‘stats’*). The age of onset of foraging, survival of bees as foragers and viral loads were all expressed relative to their respective controls (cf. Fig 1). For that purpose, individual experimental data points were subtracted from the geometric mean of their respective controls. Dunn’s Kruskal-Wallis multiple comparison tests were performed on calculated variables with Benjamini-Hochberg adjustment (package *‘FSA’*). Each treatment group was compared to its control group through Mann-Whitney tests (package *‘stats’*) on absolute values. Comparisons between experimental treatments in the immune gene expression analysis were made (-ΔΔCt) through Dunn’s tests with Benjamini-Hochberg adjustment (package *‘FSA’*). Cq results were adjusted based on the standard sample common to all plates. Adjusted Cq were used to calculate ΔCt values by subtracting each analysed sample Cq from the geometric mean of Cq for two housekeeping genes (RpS5 and RpL32). ΔΔCt values were calculated by subtracting each ΔCt in our experimental treatment from the geometric mean of ΔCt for their respective control groups. Each treatment group was compared with its respective control group through Mann-Whitney tests (package *‘stats’*) on ΔCt values.

## Results

### Behaviour and survival analysis

Among untreated bees, 47.6% (95% confidence interval (IC95) ± 4.56%) of marked bees became available foragers, suggesting that around 52.4% of them died before onset of foraging, were expelled from the hive or got lost on their first trip. Inoculation of either SBV, DWV-A or DWV-B alone did not alter the proportion of bees that became available foragers compared to their respective controls (SBV vs Sucrose: *p* = 0.54; DWVA vs PBS/Sucrose: *p* = 0.9; DWVB vs PBS/Sucrose: *p* = 0.69). We observed a significant increase in the proportion of bees that died before becoming available foragers in bees either simultaneously or sequentially co-inoculated with DWV-B and SBV compared with their control treatment or with either virus inoculation alone (DWVB/SBV vs. PBS/Sucrose: *p* = 0.032; DWVB+SBV vs. PBS+Sucrose: *p* = 1.17•10^-4^; DWVB/SBV vs. DWVB: *p* = 0.0095; DWVB+SBV vs. DWVB: *p* = 4.84•10^-6^; DWVB/SBV vs SBV: p = 3.01•10^-11^; DWVB+SBV vs SBV: p = 1.99•10^-19^ DWVB/SBV vs. PBS/SBV: *p* = 4.42•10^-24^; DWVB+SBV vs. PBS+SBV: *p* = 1.77•10^-12^; Fig 2). Surprisingly, sequential co-inoculation of PBS and SBV led to a significant decrease in the proportion of bees that died before becoming available foragers compared with its control group (*p* = 0.0194; Fig 2). While co-inoculations of DWV-A and SBV, either sequentially or simultaneously, did not alter the proportion of bees that became available foragers when compared to their respective control groups (DWVA/SBV vs PBS/Sucrose: p = 0.12; DWVA+SBV vs PBS+Sucrose: p = 0.3), significantly more mortality was observed compared to bees co-inoculated with SBV and PBS (DWVA/SBV vs PBS/SBV: p = 5.89•10^-8^; DWVA+SBV vs PBS+SBV: p = 1.34•10^-5^;  Fig 2).

**Figure 2.**
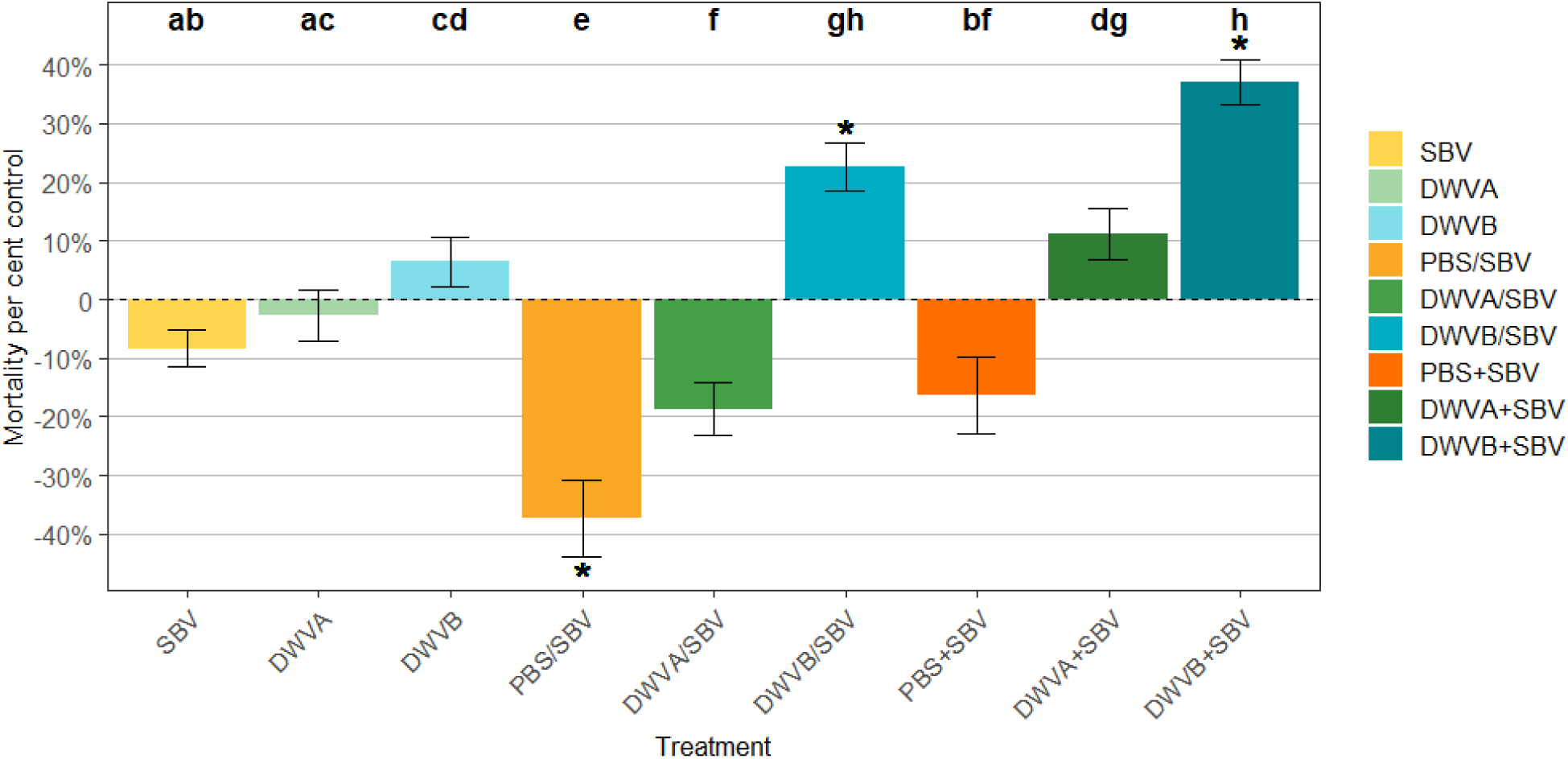
Proportions of marked bees that died before reaching their onset of foraging per cent control bees that reached their onset of foraging. Letters represent statistically different groups (p < 0.05; pairwise nominal independence test). A “*” symbol marks a statistical difference between the treated group and its control group (p < 0.05; chi-squared tests on absolute proportions). Error bars represent confidence intervals ± 95% calculated from measured values. A “/” symbol represents sequential manipulations over two different days while a “+” symbol represents simultaneous manipulations on the same day.

The next analysis focused solely on effective foragers. Untreated bees performed their first foraging trip 15.3 days (IC95 ± 1.1 days) after emergence, whereas bees injected with PBS started foraging 10.3 days (IC95 ± 0.78 days; untreated vs. PBS/Sucrose: *p* = 1.36•10^-20^; untreated vs. PBS+Sucrose: *p* = 6.84•10^-19^) after emergence. Bees inoculated with DWV-B (either alone or in co-inoculation) became foragers even earlier than control bees injected with PBS (DWVB vs. PBS/Sucrose: between 2 and 3 days earlier, *p* = 9.48•10^-8^; DWVB/SBV vs. PBS/Sucrose: between 2 and 3 days earlier, *p* = 1.75•10^-8^ and DWVB+SBV vs. PBS+Sucrose: between 3 and 4 days earlier, *p* = 1.13•10^-11^; Fig 3, A) or bees co-inoculated with PBS and SBV (DWVB/SBV vs. PBS/SBV: *p* = 4.37•10^-12^; DWVB+SBV vs. PBS+SBV: *p* = 1.92•10^-10^; Fig 3, A). Furthermore, bees simultaneously co-inoculated with DWV-B and SBV became foragers earlier than bees inoculated with DWV-B alone (between 0.5 and 1 day earlier, *p* = 0.022). Surprisingly, inoculation of DWV-A alone and sequential co-inoculation of PBS and SBV led to a slightly delayed onset of foraging compared with their control group (between 0.5 and 1 day later, *p* = 0.045 and *p* = 4.9•10^-4^, respectively; Fig 3, A). Perhaps more interestingly, we observed that, regardless of the time frame of inoculations, bees co-inoculated with DWV-A and SBV became foragers significantly earlier than bees inoculated with either DWV-A alone or co-inoculated with PBS and SBV (between 1.5 and 2 days earlier, DWVA/SBV vs. DWVA: *p* = 0.0036; DWVA/SBV vs. PBS/SBV: *p* = 7.7•10^-5^; DWVA+SBV vs. DWVA: *p* = 2.2•10^-4^; DWVA+SBV vs. PBS+SBV: *p* = 0.012; Fig 3, A).

**Figure 3.**
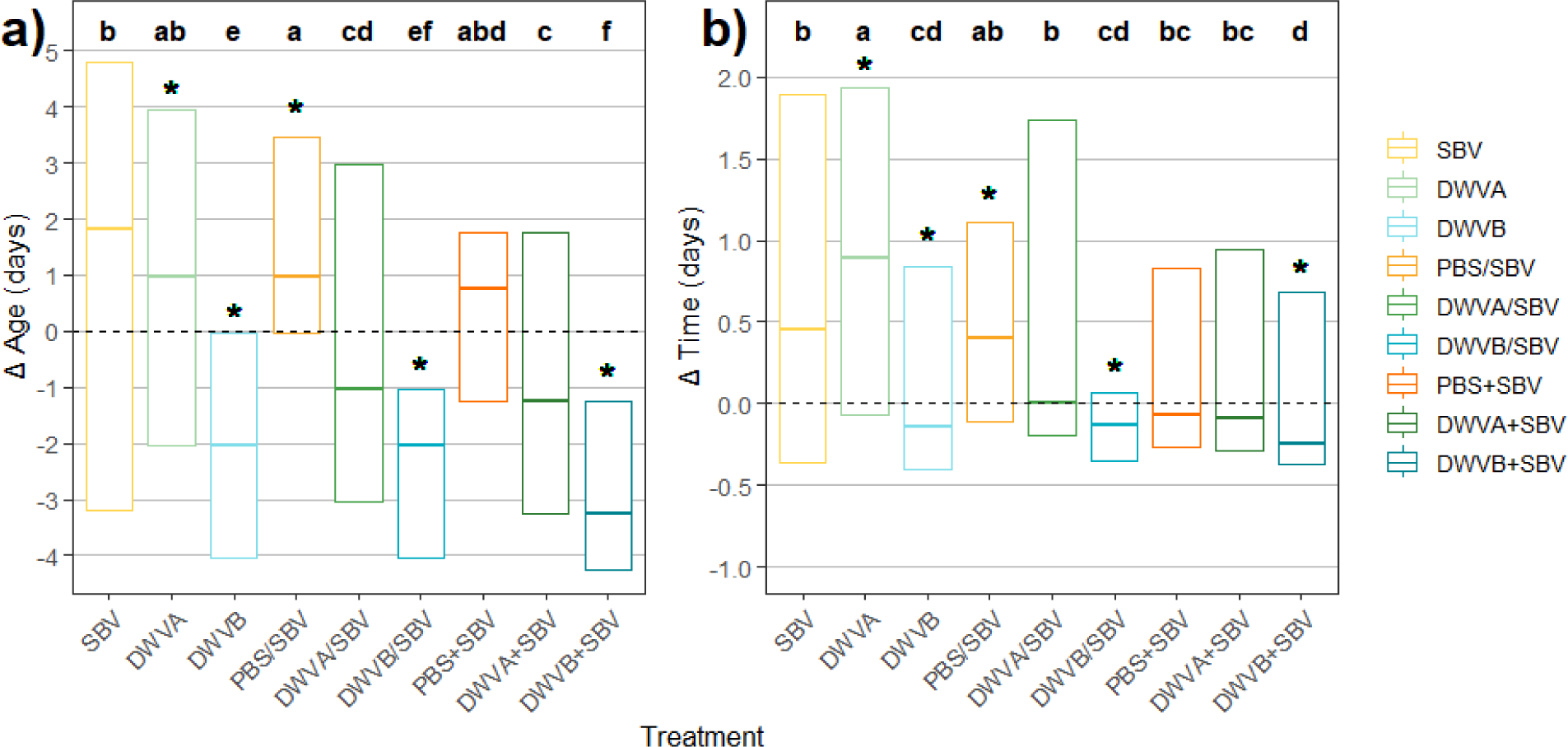
Effect of experimental treatments on the age or foragers at their onset of foraging (A) and the time between their onset of foraging and death (B), both relative to their respective control groups (cf. Fig 1). For readability purposes, whiskers were removed from boxplots, which all exceeded the shown scale. Letters represent statistically different groups (Dunn’s test). A “*” symbol marks a statistical difference between the treated group and its control group (Wilcoxon tests on absolute times). A “/” symbol represents sequential manipulations over two different days while a “+” symbol represents simultaneous manipulations on the same day.

Following their onset of foraging, untreated bees foraged for 3.9 days on average (CI95 ± 0.46 days) before dying. Control PBS injections did not significantly alter bee survival after they began foraging (untreated vs. PBS/Sucrose: *p* = 0.059; untreated vs. PBS+Sucrose: *p* = 0.091). Conversely, foragers inoculated with DWV-B (either alone or in co-inoculation with SBV) survived for significantly less time than their controls (around 0.3 days, DWVB vs. PBS/Sucrose: *p* = 0.004; DWVB/SBV vs. PBS/Sucrose: *p* = 0.011; DWVB+SBV vs. PBS+SBV: *p* = 0.0079; Fig 3, B), but co-inoculations with SBV had no effect on the survival of foragers compared with DWV-B inoculation alone (DWVB vs. DWVB/SBV: *p* = 0.9; DWVB vs. DWVB+SBV: *p* = 0.49).

### Overall virus loads

First, we analysed the frequency of viral loads for the six main honey bee viruses. SBV, DWV-A and DWV-B load frequencies followed a bimodal distribution. For SBV, most bees were found in a first peak between 10^7^ and 10^9^ genome copies per bee, while a substantial second peak was observed between 10^12^ and 10^13^ genome copies per bee. For DWV-A, most bees were found in a first peak between 10^6^ and 10^9^ genome copies per bee, while a substantial second peak was observed between 10^13^ and 10^15^ genome copies per bee. For DWV-B, most bees were found in a first peak between 10^7^ and 10^10^ genome copies per bee, while a substantial second peak was observed between 10^12^ and 10^15^ genome copies per bee. Conversely, a unimodal but not Gaussian distribution (Shapiro-Wilk tests: *p* < 0.05) was observed for ABPV, CBPV and BQCV load frequencies.

We then compared viral loads detected for DWV-A, DWV-B and SBV between each treatment, regardless of sample dates. Untreated bees were found to carry 7.2•10^7^ DWV-A genome copies (CI95 ± 0.39 log), 4.15•10^8^ DWV-B genome copies (CI95 ± 0.42 log) and 1.11•10^9^ SBV genome copies (CI95 ± 0.48 log). Control PBS and/or sucrose inoculations did not significantly alter viral loads compared with untreated bees (*p* > 0.1). As expected, inoculation of DWV-B or SBV alone led to increased loads for the respective virus compared with their respective control group (*p* = 5.38•10^-13^ and *p* = 8.76•10^-10^, respectively), although injection of DWV-B alone led to a more spectacular increase in DWV-B (of between 10^4^ and 10^5^; Fig 4, B) than the observed increase in SBV following SBV oral inoculation (of around 10^1^; Fig 4, C). However, injection of DWV-A alone did not result in a significant increase in DWV-A compared with its control group (*p* = 0.067; Fig 4, A).

**Figure 4.**
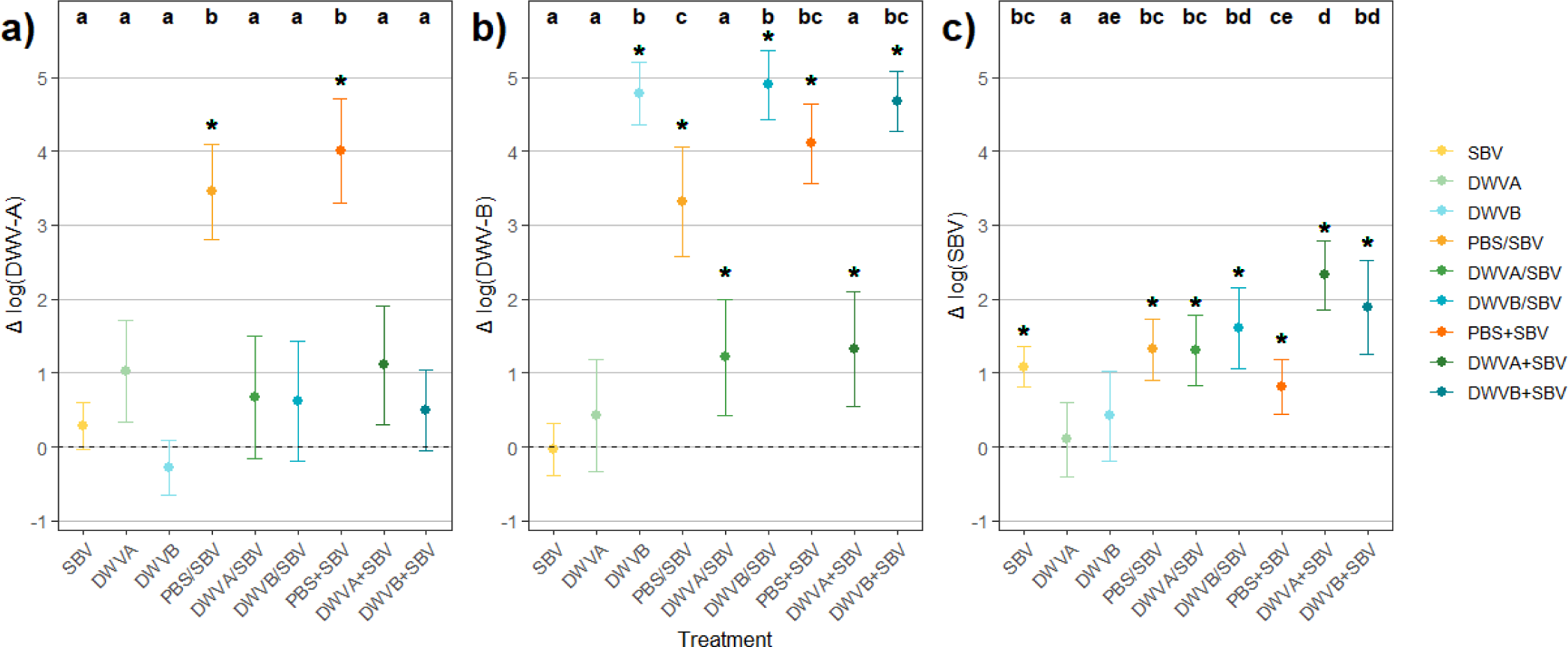
DWV-A (A), DWV-B (B) or SBV (C) loads for experimental treatments relative to their respective control groups (cf. Fig 1), irrespective of sample date. Error bars represent confidence intervals ± 95%. Letters represent statistically different groups (Dunn test). A “*” symbol marks a statistical difference between the treated group and its control group (Wilcoxon tests on absolute viral loads). A “/” symbol represents sequential manipulations over two different days while a “+” symbol represents simultaneous manipulations on the same day.

Unexpectedly, co-inoculation of both SBV and PBS led to a significant increase in both DWV genotypes compared with SBV inoculation alone or their control groups (*p* < 1•10^-9^; Fig 4, A-B). In contrast, increases in DWV-A were not observed in groups co-inoculated with either DWV-A or DWV-B and SBV regardless of the inoculations’ time frame (*p* < 0.1; Fig 4, A). Although co-inoculation with DWV-A and SBV led to significant increases in DWV-B loads compared with their control groups (DWVA/SBV vs. PBS/Sucrose: *p* = 0.045; DWVA+SBV vs. PBS+Sucrose: *p* = 0.029; Fig 4, B), DWV-B increases in bees co-inoculated with PBS and SBV were significantly higher (PBS/SBV vs. DWVA/SBV: *p* = 3.15•10^-4^; PBS+SBV vs. DWVA+SBV: *p* = 3.3•10^-6^; Fig 4, B). Increases in DWV-B among bees inoculated with DWV-B alone or co-inoculated with DWV-B and SBV were similar (DWVB vs. DWVB/SBV: *p* = 0.88; DWVB vs. DWVB+SBV: *p* = 0.86; Fig 4, B), suggesting that DWV-B alone could be responsible for these high levels of DWV-B in co-inoculation groups.

Interestingly, while inoculation of DWV-A or DWV-B alone did not result in a change in SBV loads compared with their control groups (*p* = 0.59 and *p* = 0.55, respectively), simultaneous co-inoculation with either DWV genotype and SBV led to significantly higher SBV loads than simultaneous co-inoculation of PBS and SBV (DWVA+SBV vs. PBS+SBV: *p* = 1.56•10^-4^; DWVB+SBV vs. PBS+SBV: *p* = 0.0075; Fig 4, C). Conversely, SBV loads detected in bees sequentially co-inoculated with either DWV and SBV were not statistically different from those sequentially co-inoculated with PBS and SBV (DWVA/SBV vs. PBS/SBV: *p* = 0.62; DWVB/SBV vs. PBS/SBV: *p* = 0.43; Fig 4, C).

### Viral dynamics over time

As our experimental design included two different times of inoculation (D0 and D2) and two different sampling times (D3 and D6), we were able to analyse viral dynamics over four different times post-inoculation (1 dpi, 3 dpi, 4 dpi and 6 dpi; Fig 1). However, because our experimental protocol included honey bees that endured different manipulations at different times (i.e. the sequentially inoculated groups: PBS/Sucrose, PBS/SBV, DWVA/SBV and DWVB/SBV), choices had to be made as to which inoculation we would refer to in our analysis (D0 or D2). Here, for the cited ambiguous conditions, we chose to refer to any injection when analysing DWV-A and DWV-B dynamics, and to refer to any feeding when analysing SBV dynamics. Interchanging the reference inoculations in these analyses equates to interchanging the 1 dpi results with the 3 dpi results on the one hand, and the 4 dpi with the 6 dpi results on the other for those four conditions only (Fig 1; e.g. a bee sampled on D6 could be analysed as 6 dpi if we referred to the D0 inoculation or as 4 dpi if we referred to the D2 inoculation).

The temporal analysis was used to refine the impact of inoculation chronology and virus replication dynamics. First, we observed that inoculations of DWV-B, either alone or in co-inoculation, as well as co-simultaneous co-inoculation with PBS and SBV, led to higher DWV-B loads than their controls as early as 1 dpi (DWVB vs. PBS/Sucrose: *p* = 1.97•10^-4^; DWVB/SBV vs. PBS/Sucrose: *p* = 1.17•10^-4^; DWVB+SBV vs. PBS+Sucrose: *p* = 9.79•10^-7^; PBS+SBV vs. PBS+Sucrose: *p* = 7.29•10^-5^) and remained at higher levels than their controls on 3 dpi (*p* = 4.6•10^-5^, *p* = 4.6•10^-5^, *p* = 1.4•10^-7^ and *p* = 3.45•10^-4^, respectively), 4 dpi (*p* = 6.29•10^-6^, *p* = 4.47•10^-5^, *p* = 2.57•10^-6^ and *p* = 3.54•10^-8^, respectively) and 6 dpi (*p* = 3.76•10^-5^, *p* = 3.48•10^-5^, *p* = 4.5•10^-6^ and *p* = 2.94•10^-5^, respectively; Fig 5, B). Conversely, sequential co-inoculation of PBS and SBV did not lead to significantly elevated SBV loads on 1 dpi (*p* = 0.096), but did reach significantly higher DWV-B loads on 3 dpi (*p* = 2.16•10^-5^), 4 dpi (*p* = 0.022) and 6 dpi (*p* = 1.38•10^-5^) than its control group (Fig 5, B). The previous observation that DWV-A and SBV co-inoculation led to increases in DWV-B loads was due to significant increases in DWV-B loads on 4 dpi only (DWVA/SBV vs. PBS/Sucrose: *p* = 0.043; DWVA+SBV vs. PBS+Sucrose: *p* = 0.0088), but DWV-B levels for the two groups were again similar to their respective controls on 6 dpi (DWVA/SBV vs. PBS/Sucrose: *p* = 0.47; DWVA+SBV vs. PBS+Sucrose: *p* = 74).

**Figure 5.**
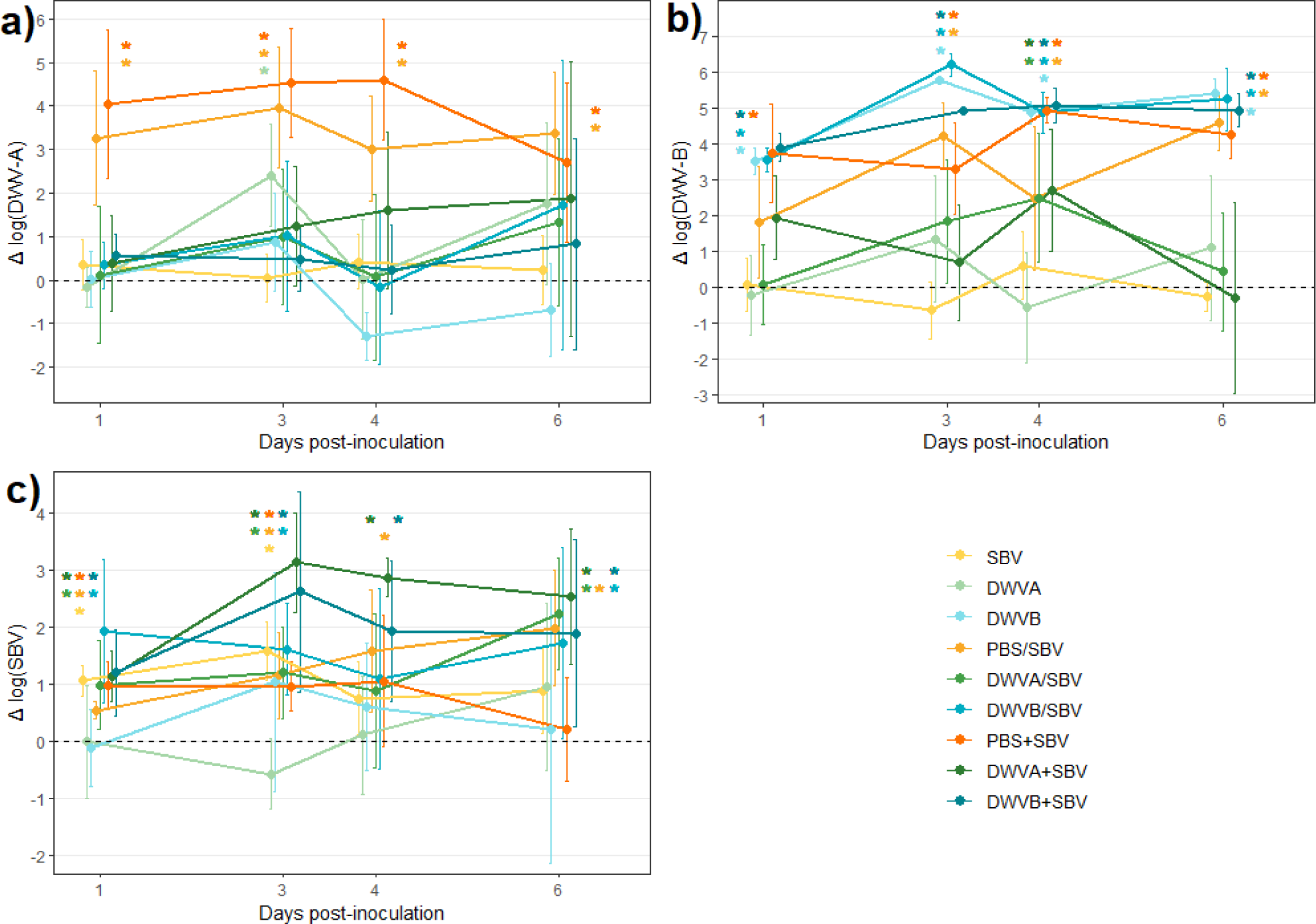
Evolution of DWV-A (A), DWV-B (B) or SBV (C) loads over time relative to their respective control groups. Error bars represent confidence intervals ± 95%. A “*” symbol represents a statistical difference between the treated group and its control group on 6 dpi (Wilcoxon tests on absolute viral loads). A “/” symbol represents sequential manipulations over two different days while a “+” symbol represents simultaneous manipulations on the same day.

Bees subjected to either sequential or simultaneous co-inoculation with PBS and SBV also carried higher DWV-A loads than their respective controls as early as 1 dpi (*p* = 1.35•10^-4^ and *p* = 1.35•10^-4^, respectively) and remained higher than their controls on 3 dpi (*p* = 3•10^-5^ and *p* = 1.1•10^-6^, respectively), 4 dpi (*p* = 0.0041 and *p* = 4•10^-5^, respectively) and 6 dpi (*p* = 1.73•10^-4^ and *p* = 0.0072, respectively; Fig 5, A).

Surprisingly, different SBV dynamics were observed depending on whether PBS and SBV were inoculated sequentially or simultaneously. While bees inoculated with SBV alone and bees co-inoculated either simultaneously or sequentially with PBS and SBV all showed significantly elevated SBV loads compared with their control groups on 1 dpi (SBV vs. Sucrose: *p* = 2.07•10^-5^; PBS/SBV vs. PBS/Sucrose: *p* = 0.0054; PBS+SBV vs. PBS+Sucrose: *p* = 0.024) and 3 dpi (SBV vs. Sucrose: *p* = 2.34•10^-7^; PBS/SBV vs. PBS/Sucrose: *p* = 0.016; PBS+SBV vs. PBS+Sucrose: *p* = 0.0018), this was only the case for bees sequentially co-inoculated with PBS and SBV on 4 dpi (SBV vs. Sucrose: *p* = 0.069; PBS/SBV vs. PBS/Sucrose: *p* = 0.0036; PBS+SBV vs. PBS+Sucrose: *p* = 0.13) and 6 dpi (SBV vs. Sucrose: *p* = 0.073; PBS/SBV vs. PBS/Sucrose: *p* = 0.0048; PBS+SBV vs. PBS+Sucrose: *p* = 0.035; Fig 5, C). Compared with their control groups, we also observed significantly higher SBV loads in bees either simultaneously or sequentially co-inoculated with either DWV-A or DWV-B and SBV as early as 1 dpi (DWVA/SBV vs. PBS/Sucrose: *p* = 0.0036; DWVB/SBV vs. PBS/Sucrose: *p* = 0.0023; DWVA+SBV vs. PBS+Sucrose: *p* = 0.013; DWVB+SBV vs. PBS+Sucrose: *p* = 0.025), which remained higher than their respective controls on 3 dpi (*p* = 2.1•10^-6^ and *p* = 0.0021, respectively), 4 dpi (*p* = 0.0023 and *p* = 0.033, respectively) and 6 dpi (*p* = 0.013 and *p* = 0.036, respectively) for simultaneous co-inoculation and on 3 dpi (*p* = 0.0093 and *p* = 0.015, respectively) and 6 dpi (*p* = 0.0018 and *p* = 0.021, respectively) for sequential co-inoculation (Fig 5, C).

### Immune gene expression

In this study, we had the opportunity to analyse the immune profile of bees in different states of infection: SBV inoculation alone led to high SBV loads only; DWV-B inoculation alone led to high DWV-B loads only; DWV-A inoculation alone led to moderately high DWV-A loads, co-inoculation of DWV-A and SBV led to high DWV-A and SBV loads; co-inoculation of DWV-B and SBV led to high DWV-B and SBV loads; co-inoculation of PBS and SBV led to high DWV-A, DWV-B and SBV loads. This section gives the results of the analysed expression of eight immune genes involved in different immune pathways (Fig 6).

**Figure 6.**
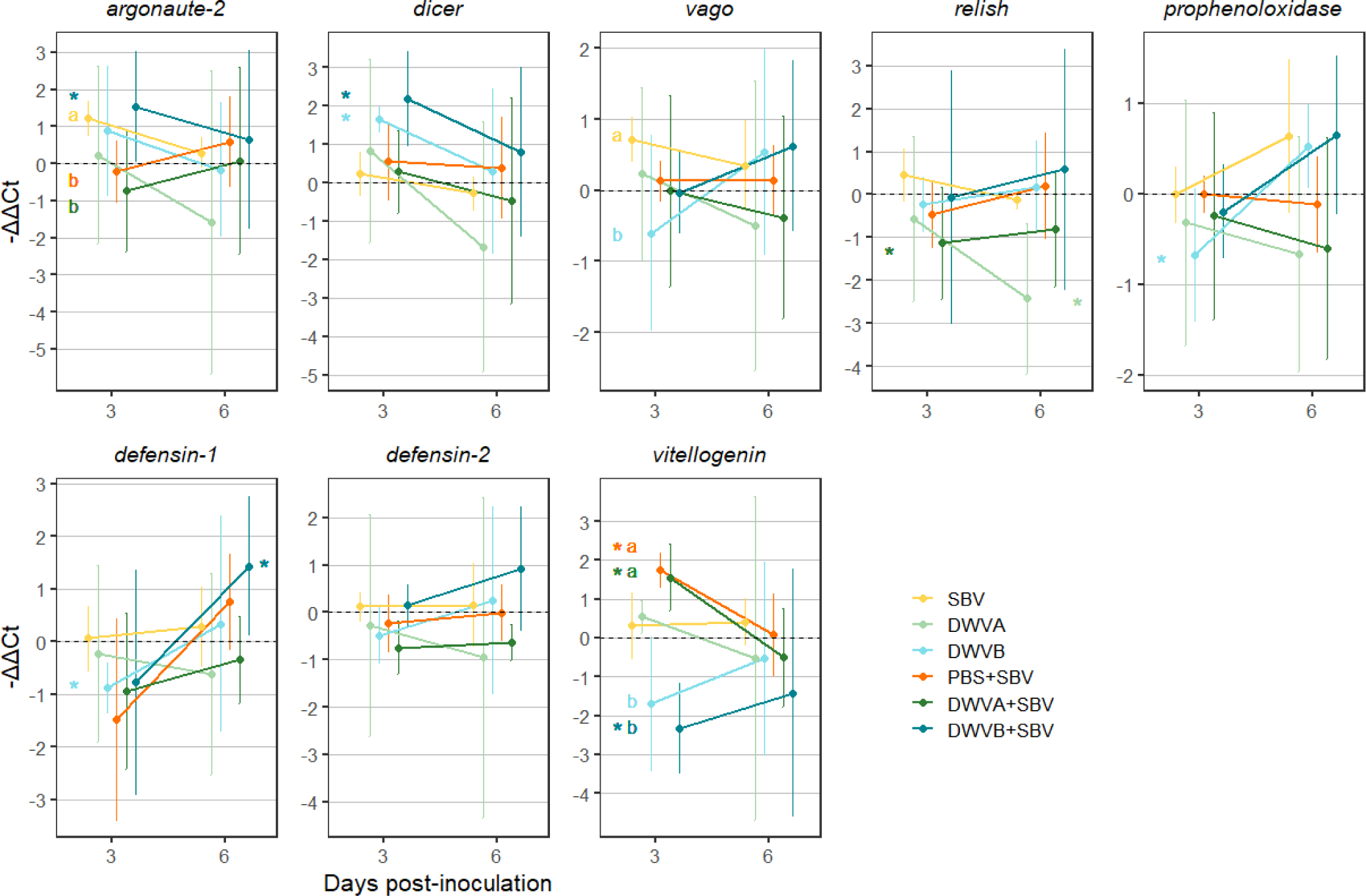
Expression of immune genes on two different days post-inoculation. Gene expression is expressed as (-ΔΔCt). Error bars represent confidence intervals ± 95%. Letters represent statistical differences between experimental groups (p < 0.05; Dunn tests on (-ΔΔCt)). A “*” symbol represents a significant difference between an experimental group and its control group (p < 0.05; Wilcoxon tests on ΔCt values).

Compared with their respective control group, DWV-B inoculation alone triggered a higher *dicer* expression (*p* = 0.024), and lower *prophenoloxidase* and *defensin-1* expression on 3 dpi (*p* = 0.048 and *p* = 0.024, respectively). Co-inoculation with DWV-B and SBV led to a higher expression of *argonaute-2* and *dicer* on 3 dpi (*p* = 0.024 for both comparisons), lower expression of *vitellogenin* on 3 dpi (*p* = 0.024), and higher expression of *defensin-1* on 6 dpi (*p* = 0.048). Inoculation of DWV-A alone only led to a decreased expression of *relish* on 6 dpi (*p* = 0.024) while co-inoculation with DWV-A and SBV led to decreased expression of *relish* on 3 dpi (*p* = 0.024) and increased expression of *vitellogenin* on 3 dpi (*p* = 0.024). Co-inoculation with PBS and SBV only led to an increased expression of *vitellogenin* on 3 dpi (*p* = 0.0022). Bees inoculated with SBV alone expressed significantly more *argonaute-2* on 3 dpi than bees co-inoculated with DWV-A and SBV or co-inoculated with PBS and SBV (*p* = 0.038 and *p* = 0.047, respectively), and significantly more *vago* on 3 dpi than bees inoculated with DWV-B alone (*p* = 0.021). Finally, bees co-inoculated with DWV-A and SBV, and bees co-inoculated with PBS and SBV expressed significantly more *vitellogenin* on 3 dpi than bees inoculated with DWV-B, whether alone or in co-inoculation (PBS+SBV vs. DWVB: *p* = 0.012; PBS+SBV vs. DWVB+SBV: *p* = 0.007; DWVA+SBV vs. DWVB: *p* = 0.03; DWVA+SBV vs. DWVB+SBV: *p* = 0.016).

## Discussion

To our knowledge, no previous studies have investigated virus co-inoculation in adult honey bees. Here, we show that inoculating bees with DWV-B reduces the survival of foragers, and that its co-inoculation with SBV synergistically amplifies DWV-B – but not DWV-A – virulence in adult bees. This synergistical effect was found in the mortality rates of nurse bees as well as in the precocious onset of foraging of bees. We also found three distinct effects of DWV and SBV co-infection on viral dynamics. Firstly, co-inoculation with either DWV genotype and SBV synergistically promotes SBV replication. Secondly, inoculation of SBV combined with an injection-like injury drastically promotes the replication of DWV already present in bees. But thirdly, this effect was not observed when superinfecting DWV was co-inoculated, suggesting competition between the DWV already present in the bee and the superinfecting DWV. Finally, we found that co-inoculations often led to intensified immune responses compared to DWV inoculations alone, and that inoculation of different DWV genotypes can lead to opposite responses.

The viral loads and distributions of ABPV (mean load found at 4.92 log IC95 ± 0.16 log), CBPV (mean load found at 5.47 log IC95 ± 0.06 log) and BQCV (mean load found at 7.0 log IC95 ± 0.11 log) viral load frequencies found in this study suggest that these three viruses did not interfere with our results. Indeed in naturally infected bees, ABPV and CBPV viral load frequencies follow a bimodal distribution [50]. In this study, we found unimodal distributions of these viral load frequencies (S2 Fig). Given the high virulence of both paralysis bee viruses, we can reasonably assume that virtually none of the analysed bees were overtly infected by either of these two viruses. In the case of BQCV, our results are consistent with the literature, with a wide prevalence of the virus with no observation of two distinct populations among infected bees [50].

In this study, we observed that co-inoculation with either DWV genotype and SBV tends to further promote SBV replication compared with virus inoculations alone. Such an increase in SBV replication was previously found in pupae injected with DWV (Dubois et al., 2020). While this increased SBV replication may not be pathogenic for individual adult bees and would not increase transmission between older adults [53], it may promote increased SBV transmission between nurse bees and larvae, thus increasing larvae mortality and further weakening the colony. However, in bees sequentially co-inoculated with DWV-B and SBV, the observed increase in SBV was only observed from 6 days post-inoculation, an age from which these bees start to become foragers. The early onset of foraging of these bees may compensate for the potential increase in SBV transmission to larvae [54]. To address both those hypotheses, the potential of SBV to be transmitted to larvae from differentially infected adult bees, either directly from nurse bees or indirectly through the nectar collected by foragers, should be investigated in future studies.

In this study, we observed that DWV-A and SBV inoculated alone did not negatively impact honey bee survival nor induce precocious onset of foraging. Conversely, DWV-B inoculated alone did not negatively impact honey bee survival before their onset of foraging (living time spent as nurse bees), but induced an early onset of foraging and decreased the survival of foragers. Here, we showed that either sequential or simultaneous co-inoculation of DWV-B and SBV synergistically increased the mortality of nurse bees and simultaneous co-inoculations precipitated the onset of survivors’ foraging. The decrease in survival of bees that became foragers was, however, similar between bees inoculated with DWV-B alone or co-inoculated with DWV-B and SBV. Thus, through increased mortality in nurse bees and a more precocious onset of foraging, simultaneous co-exposure of bees to DWV-B and SBV may significantly reduce the nurse workforce while also decreasing the pool of available foragers. Furthermore, co-inoculation did not mitigate the negative impact of DWV-B inoculated alone on the survival of foragers. This may have dire consequences on the colony’s capacity to forage and store resources as well as on its ability to properly carry out in-hive duties such as brood care. Combined with the direct consequences of increased mortality within the colony, co-exposure of bees to DWV-B and SBV may further weaken the colony compared with exposure to one or other of the viruses. One study [55] compared DWV, SBV and BQCV loads and overwinter mortality in varroa-sensible (MS) and varroa-resistant (MR) colonies. They found better colony survival in MR colonies but found similar DWV loads in MR and MS colonies. However, SBV and BQCV loads were significantly lower in MR colonies than in MS colonies. This is in line with our own results, and suggests that the increased virulence of DWV in the presence of SBV may indeed have tangible consequences for colony health and survival.

In our experiments, co-inoculation occurred either within a 2-hour time frame (simultaneous co-inoculation) or 48 hours apart (sequential co-inoculation). Our results suggest that the time between inoculations plays a role in the emergence of synergies between DWV-B and SBV. While both co-inoculation time-frames led to a synergistical decrease in available foragers for the colony (Fig 2), only the simultaneous co-inoculation further decreased the age of onset of foraging of bees compared to DWV-B inoculation alone (Fig 3). While leading to similar levels in SBV loads on 6 days post-inoculation, we observed different dynamics in bees either simultaneously or sequentially co-inoculated (Fig 5), with a more stable increase in SBV loads for simultaneous co-inoculation groups. Interestingly, in bees subjected to SBV inoculation combined with an injection injury (PBS injection), stable increases in SBV loads were only observed in the sequential inoculation scenario. In contrast, the dynamics for both DWV-A and DWV-B loads were similar between the sequential and simultaneous scenario in these bees. Therefore, we can infer that replication of DWV genotypes that were already present may not promote SBV replication, that superinfecting DWV may specifically promote SBV replication when simultaneously co-inoculated, and that cuticular injury chronologically distant from SBV exposure may further promote SBV replication independently of DWV. Overall, while sequential co-inoculation had a tendency to drive synergies between DWV and SBV, simultaneous co-inoculation revealed them more clearly.

Compared with our experimental setting, bees in natural conditions are more likely to acquire lower quantities of DWV and SBV. However, they are also more likely to be exposed to both viruses way more often. On the one hand, SBV is a highly prevalent virus among bees [34]. Social interactions occur frequently within the colony, as does removal of diseased larvae and pupae together with cannibalism [56,57]. This combination may lead to recurrent exposure of bees to SBV. On the other hand, our injection method aimed to mimic exposure of bees to varroa-mediated injury and DWV transmission. While little is known about the frequency of varroa bites, a recent study found that varroa mites regularly switch hosts, some of them even changing daily (Lamas et al., 2023). This would suggest that both simultaneous and sequential co-inoculations of SBV and DWV (especially the DWV-B genotype) could occur regularly in natural conditions. Furthermore, our results suggest that bites from varroa mites free of DWV could still trigger viral replication in honey bees, as we showed that exposure to SBV combined with an injection-like injury can lead to increased replication of background DWV genotypes already present in bees. These increases in background DWV loads were even higher when DWV was not inoculated, suggesting potential competition between DWV already present and some superinfecting DWV genotypes. This competition may still be conditional, as superinfecting DWV-B still triggered a high increase in DWV-B when co-inoculated with SBV. As we do not know whether the observed replication of DWV-B originated from our inoculum or background DWV-B, we cannot infer whether one DWV-B genotype outcompeted the other or if an absence of competition allowed both genotypes to replicate. Although varroa mites are known to commonly vector DWV, the viral loads of varroa mites could be subjected to seasonality, as we detected few to no DWV-B in mites sampled in November 2022 and February 2019 in our experimental apiary, but detected comparatively higher loads in mites sampled at the end of March 2019 (S2 Table). This scenario would thus most likely occur during late autumn or early winter, although further experiments could unravel the dynamics of DWV infections in varroa in greater detail. Finally, while the observed increases in DWV loads would probably not be as drastic in natural conditions than in our study, we can still expect increases in DWV replication in bees both injured by varroa mites and repeatedly exposed to SBV at lower doses.

Overall, the greatest change in immune gene expression was found in bees co-inoculated with DWV-B and SBV (a significantly higher expression of *argonaute-2* and *dicer* on 3 dpi, *defensin-1* on 6 dpi and significantly lower expression of *vitellogenin* on 3 dpi, compared with its control group). The greatest mortality rates were also found for this group of co-inoculated bees (Fig 1-2). Indeed, immune responses are known to be costly for bees [58,59]. It has also been shown that greater immune responses may be linked to higher susceptibility to pathogen replication in young bees [60]. Other authors found decreases in *vitellogenin* expression in DWV-inoculated bees [47,61]. In our study, *vitellogenin* expression was only found to be significantly decreased in bees co-inoculated with DWV-B and SBV. Immune responses require Vitellogenin [62,63], so an elevated immune response combined with decreased *vitellogenin* expression may lead to a drastic depletion of Vitellogenin stores. As Vitellogenin levels control task allocation [64] and are linked to longevity [62], the depletion of Vitellogenin stores may explain both the increased mortality observed and the early onset of foraging of these co-inoculated bees. However, we did not find the opposite to be true, as simultaneous DWV-A and SBV co-inoculation and simultaneous PBS and SBV co-inoculation led to significantly elevated *vitellogenin* expression, but no change in survival or age of the onset of foraging was observed compared with their control group. Interestingly, unlike DWV-B replication triggered by a superinfecting DWV-B, high replication of DWV-B already present in bees did not lead to either pronounced immune responses or elevated mortality.

Additionnally, interesting tendencies may be discussed. Firstly, the immune gene expression profiles of co-inoculated bees tended to be more similar to those of bees inoculated with DWV alone than to those of bees inoculated with SBV alone (e.g. *dicer* on 3 dpi, *vago* and *prophenoloxidase* on 6 dpi). Co-inoculation of SBV with one DWV genotype tended to lead to a qualitatively similar (increase or decrease in expression) but quantitatively exacerbated response compared with DWV inoculation alone (e.g. *argonaute-2*, *dicer* and *vitellogenin* on 3 dpi, *defensin-1* and *defensin-2* on 6 dpi). Secondly, co-inoculation with SBV, DWV-A and DWV-B tended to trigger opposite responses (e.g. *argonaute-2*, *dicer*, *relish* and *vitellogenin* on 3 dpi, *vago*, *prophenoloxidase*, *defensin-1* and *defensin-2* on 6 dpi). Inoculations of DWV-A, either alone or in co-inoculation, did not trigger elevated immune responses despite this being the case for DWV-B inoculations. This could either originate from the DWV genotype or the degree of viral replication, as DWV-B replicated more than DWV-A when inoculated (Fig. 4-5). However, a high replication of both DWV genotypes in bees co-inoculated with PBS and SBV led to gene expression similar to that of bees co-inoculated with DWV-A and SBV (e.g. *argonaute-2* and *vitellogenin* on 3 dpi). This might suggest that DWV-A pressure may specifically inhibit the RNAi pathway and promote *vitelogenin* expression.

When observed, the RNAi pathway was activated on 3 dpi but not on 6 dpi, confirming that this pathway is an early defence system against viruses [65]. However, other studies have found RNAi gene transcription activated as late as 5 dpi [66,67]. Similarly to our results, one study [67] found that inhibition of *defensin-1* can occur in parallel to the activation of the RNAi pathway in DWV-B inoculated bees. Our study shows that there may be concurrent inhibition of the melanisation pathway, which was not the case in other studies [40,68]. Conversely, when observed, AMP production was activated on 6 dpi but not on 3 dpi, confirming that AMPs are used as a late defence system that may remain active for longer times [61]. However, our study highlights the importance of DWV genotypes and co-inoculated viruses in the bee immune response.

Honey bee health has become a major concern in the past decades. As research has progressed, the network of interactions between honey bee stressors has been shown to be more complex than previously thought. Here, we introduce viral interactions as a significant additional level of complexity. According to our results, SBV may play a role in the increased DWV virulence and replication observed since the introduction of *Varroa destructor*. Moreover, interactions between the two viruses may also favour SBV replication, potentially leading to increased frequency of co-exposure. Our study highlights the importance of viral interactions and the need for future research to consider the complexity of the honey bee virobiome in the study of honey bee health.

## Author contributions

**Conceptualization** A.B-D., E.D., AM.D.; **Data Curation** AM.D.; **Formal Analysis** AM.D.; **Funding Acquisition** A.B-D., E.D.; **Investigation** AM.D., A.B-D.; **Methodology** AM.D., A.B-D.; **Project Administration** AM.D., A.B-D.; **Resources** A.B-D.; **Software** AM.D. **Supervision** A.B-D.; **Validation** A.B-D.; **Visualization** AM.D.; **Writing – Original Draft Preparation** AM.D.; **Writing – Review and Editing** AM.D., A.B-D., E.D.

## Acknowledgements

The authors would like to thank Didier Crauser for handling the optical counters and Émilien Rottier for managing the apiary. We also thank Virginie Diévart for managing the molecular biology laboratory. We are grateful to Joy Giraud, Bao-Huynh Nguyen, Elsa Frileux and Émilie Balthazar for their help in marking and infecting the bees, and to Dr. Vincent Piou for collecting the mites in 2019 and performing the RNA purification and cDNA synthesis on them. We thank the participants of the Bioproject (PRJNA1055031) from which our viral strains originated. Finally, the authors wish to thank Delphine Libby-Claybrough, professional translator and native English speaker, for her editorial assistance in revising the manuscript.

## Funding

This research was funded by both the National Research Institute for Agriculture Food and Environement, INRAE, and the French Agency for Food, Environmental and Occupational Health Safety, ANSES.

## Data availability statement

The DOIs for honey bee survival, behaviour, viral loads and immune genes expression datasets will be made available after acceptance of the paper.

## Supporting information

**S1 Table Virus and immune gene primers.** Primers used for virus quantification and immune gene expression analysis and their references.

**S2 Table Sample sizes for each treatment and analysis.** For each treatment, number of host colonies tested, number of bees maked and detected at each stage, number of bees analysed in virus quantification and immune gene expression analysis.

**S1 Figure Vizualisation of marked frame-bees and optical counter recording.** A) Picture of a hive frame section including painted bees, QR-code-marked bees and the painted queen. B) Picture of a bees passing through an optical counter tunnel.

**S2 Figure Viral load frequency distributions for the six quantified viruses.** Graphical representation of SBV, DWV-A, DWV-B, BQCV, ABPV and CBPV viral load frequency distributions in all analysed bees.

**S3 Table Viral loads of Varroa destructor mites collected throughout the year 2019 and 2021.** DWV-A, DWV-B, ABPV, BQCV and SBV viral quantifications performed on Varroa destructor mites collected on automn 2021, winter 2019 and spring 2019 on the apiary of ANSES Sophia Antipolis.

## Notes

### Competing Interest Statement

The authors have declared no competing interest.

## References

1. Beaurepaire A, Piot N, Doublet V, Antunez K, Campbell E, Chantawannakul P, et al. Diversity and global distribution of viruses of the western honey bee, apis mellifera. Insects. 2020;11(4):1–25.

2. Cox-Foster DL, Conlan S, Holmes EC, Palacios G, Evans JD, Moran NA, et al. A Metagenomic Survey of Microbes in Honey Bee Colony Collapse Disorder. Science (80-). 2007;318(October):283–8.

3. Dainat B, Evans JD, Chen YP, Gauthier L, Neumann P. Predictive markers of honey bee colony collapse. PLoS One. 2012;7(2).

4. Wei R, Cao L, Feng Y, Chen Y, Chen G, Zheng H. Sacbrood Virus: A Growing Threat to Honeybees and Wild Pollinators. Viruses. 2022;14(9).

5. Mazzei M, Carrozza ML, Luisi E, Forzan M, Giusti M, Sagona S, et al. Infectivity of DWV associated to flower pollen: Experimental evidence of a horizontal transmission route. PLoS One. 2014;9(11):1–16.

6. Locke B, Semberg E, Forsgren E, De Miranda JR. Persistence of subclinical deformed wing virus infections in honeybees following Varroa mite removal and a bee population turnover. PLoS One. 2017;12(7):1–10.

7. Yue C, Schröder M, Gisder S, Genersch E. Vertical-transmission routes for deformed wing virus of honeybees (Apis mellifera). J Gen Virol. 2007;88(8):2329–36.

8. de Graaf DC, Laget D, De Smet L, Claeys Boúúaert D, Brunain M, Veerkamp RF, et al. Heritability estimates of the novel trait ‘suppressed in ovo virus infection’ in honey bees (Apis mellifera). Sci Rep [Internet]. 2020;10(1):1–10. Available from: 10.1038/s41598-020-71388-x

9. de Miranda JR, Fries I. Venereal and vertical transmission of deformed wing virus in honeybees (Apis mellifera L.). J Invertebr Pathol. 2008;98(2):184–9.

10. Oldroyd BP. Coevolution while you wait: Varroa jacobsoni, a new parasite of western honeybees. Trends Ecol Evol. 1999;14(8):312–5.

11. Koeniger N, Koeniger G, Wijayagunasekara NHP. Beobachtungen über die Anpassung von Varroa jacobsoni an ihren natürlichen Wirt Apis cerana in Sri Lanka (Observations on the Adaptation of Varroa Jacobsoni to its Natural Host Apis Cerana in Sri Lanka) (in German). Apidologie. 1981;12(1):37–40.

12. Guzman LI De, Rinderer TE, Guzman LI De, Identification TER, Guzman LI De. Identification and comparison of Varroa species infesting honey bees To cite this version: HAL Id: hal-00891570 species infesting honey bees. 1999;

13. Kasangaki P, Sarah Otim A, P’Odyek Abila P, Angiro C, Chemurot M, Kajobe R. Presencia de Varroa en Uganda y su conocimiento por parte de la industria apícola. J Apic Res [Internet]. 2015;54(4):373–7. Available from: 10.1080/00218839.2016.1159858

14. Santamaria J, Villalobos EM, Brettell LE, Nikaido S, Graham JR, Martin S. Evidence of Varroa-mediated deformed wing virus spillover in Hawaii. J Invertebr Pathol [Internet]. 2018;151(November 2017):126–30. Available from: 10.1016/j.jip.2017.11.008

15. Chapman NC, Colin T, Cook J, Da Silva CRB, Gloag R, Hogendoorn K, et al. The final frontier: Ecological and evolutionary dynamics of a global parasite invasion. Biol Lett. 2023;19(5).

16. Annoscia D, Brown SP, Di Prisco G, De Paoli E, Del Fabbro S, Frizzera D, et al. Haemolymph removal by Varroa mite destabilizes the dynamical interaction between immune effectors and virus in bees, as predicted by Volterra’s model. Proc R Soc B Biol Sci. 2019;286(1901).

17. Ramsey SD, Ochoa R, Bauchan G, Gulbronson C, Mowery JD, Cohen A, et al. Varroa destructor feeds primarily on honey bee fat body tissue and not hemolymph. Proc Natl Acad Sci U S A. 2019;116(5):1792–801.

18. Di Prisco G, Pennacchio F, Caprio E, Boncristiani HF, Evans JD, Chen Y. Varroa destructor is an effective vector of Israeli acute paralysis virus in the honeybee, Apis mellifera. J Gen Virol. 2011;92(1):151–5.

19. Möckel N, Gisder S, Genersch E. Horizontal transmission of deformed wing virus: Pathological consequences in adult bees (Apis mellifera) depend on the transmission route. J Gen Virol. 2011;92(2):370–7.

20. Locke B, Forsgren E, Fries I, de Miranda JR. Acaricide treatment affects viral dynamics in Varroa destructor-infested honey bee colonies via both host physiology and mite control. Appl Environ Microbiol. 2012;78(1):227–35.

21. Posada-Florez F, Ryabov E V., Heerman MC, Chen Y, Evans JD, Sonenshine DE, et al. Varroa destructor mites vector and transmit pathogenic honey bee viruses acquired from an artificial diet. PLoS One [Internet]. 2020;15(11 November):1–13. Available from: 10.1371/journal.pone.0242688

22. Gisder S, Aumeier P, Genersch E. Deformed wing virus: Replication and viral load in mites (Varroa destructor). J Gen Virol. 2009;90(2):463–7.

23. Gisder S, Genersch E. Direct Evidence for Infection of Varroa destructor Mites with the Bee-Pathogenic Deformed Wing Virus Variant B, but Not Variant A, via Fluorescence In Situ Hybridization Analysis. J Virol. 2021;95(5).

24. Martin SJ, Highfield AC, Brettell L, Villalobos EM, Budge GE, Powell M, et al. Global honey bee viral landscape altered by a parasitic mite. Science (80-). 2012;336(6086):1304–6.

25. Ray AM, Davis SL, Rasgon JL, Grozinger CM. Simulated vector transmission differentially influences dynamics of two viral variants of deformed wing virus in honey bees (Apis mellifera). J Gen Virol. 2021;102(11).

26. Ryabov E V., Childers AK, Lopez D, Grubbs K, Posada-Florez F, Weaver D, et al. Dynamic evolution in the key honey bee pathogen deformed wing virus: Novel insights into virulence and competition using reverse genetics. PLoS Biol [Internet]. 2019;17(10):1–27. Available from: 10.1371/journal.pbio.3000502

27. Moore J, Jironkin A, Chandler D, Burroughs N, Evans DJ, Ryabov E V. Recombinants between Deformed wing virus and Varroa destructor virus-1 may prevail in Varroa destructor-infested honeybee colonies. J Gen Virol. 2011;92(1):156–61.

28. Dalmon A, Desbiez C, Coulon M, Thomasson M, Le Conte Y, Alaux C, et al. Evidence for positive selection and recombination hotspots in Deformed wing virus (DWV). Sci Rep [Internet]. 2017;7(June 2016):1–12. Available from: 10.1038/srep41045

29. McMahon DP, Natsopoulou ME, Doublet V, Fürst M, Weging S, Brown MJF, et al. Elevated virulence of an emerging viral genotype as a driver of honeybee loss. Proc R Soc B Biol Sci. 2016;283(1833).

30. Natsopoulou ME, McMahon DP, Doublet V, Frey E, Rosenkranz P, Paxton RJ. The virulent, emerging genotype B of Deformed wing virus is closely linked to overwinter honeybee worker loss. Sci Rep. 2017;7(1):1–9.

31. Norton AM, Remnant EJ, Buchmann G, Beekman M. Accumulation and Competition Amongst Deformed Wing Virus Genotypes in Naïve Australian Honeybees Provides Insight Into the Increasing Global Prevalence of Genotype B. Front Microbiol. 2020;11(April):1–14.

32. Paxton RJ, Schäfer MO, Nazzi F, Zanni V, Annoscia D, Marroni F, et al. Epidemiology of a major honey bee pathogen, deformed wing virus: potential worldwide replacement of genotype A by genotype B. Int J Parasitol Parasites Wildl. 2022;18(March):157–71.

33. Doublet V, Oddie MAY, Mondet F, Forsgren E, Dahle B, Furuseth-Hansen E, et al. Shift in virus composition in honeybees (Apis mellifera) following worldwide invasion by the parasitic mite and virus vector Varroa destructor. R Soc Open Sci. 2024;11(1).

34. Tentcheva D, Gauthier L, Zappulla N, Dainat B, Cousserans F, Colin ME, et al. Prevalence and seasonal variations of six bee viruses in Apis mellifera L. and Varroa destructor mite populations in France. Appl Environ Microbiol. 2004;70(12):7185–91.

35. Roy C, Vidal-Naquet N, Provost B. A severe sacbrood virus outbreak in a honeybee (Apis mellifera L.) colony: A case report. Vet Med (Praha) [Internet]. 2015;60(6):330–5. Available from: 10.17221/8248-VETMED

36. Han B, Zhang L, Feng M, Fang Y, Li J. An integrated proteomics reveals pathological mechanism of honeybee (Apis cerena) sacbrood disease. J Proteome Res. 2013;12(4):1881–97.

37. Shan L, Liuhao W, Jun G, Yujie T, Yanping C, Jie W, et al. Chinese Sacbrood virus infection in Asian honey bees (Apis cerana cerana) and host immune responses to the virus infection. J Invertebr Pathol [Internet]. 2017;150(September):63–9. Available from: 10.1016/j.jip.2017.09.006

38. Chang JC, Chang ZT, Ko CY, Scotty Yang CC, Chen YW, Nai YS. Sacbrood viruses cross-infection between Apis cerana and Apis mellifera: Rapid detection, viral dynamics, evolution and spillover risk assessment. J Invertebr Pathol [Internet]. 2021;186(October):107687. Available from: 10.1016/j.jip.2021.107687

39. Duffy S. Why are RNA virus mutation rates so damn high? PLoS Biol. 2018;16(8):1–6.

40. Ryabov E V., Fannon JM, Moore JD, Wood GR, Evans DJ. The Iflaviruses Sacbrood virus and Deformed wing virus evoke different transcriptional responses in the honeybee which may facilitate their horizontal or vertical transmission. PeerJ. 2016;2016(1).

41. Remnant EJ, Mather N, Gillard TL, Yagound B, Beekman M. Direct transmission by injection affects competition among RNA viruses in honeybees. Proc R Soc B Biol Sci. 2019;286(1895).

42. Mondet F, de Miranda JR, Kretzschmar A, Le Conte Y, Mercer AR. On the Front Line: Quantitative Virus Dynamics in Honeybee (Apis mellifera L.) Colonies along a New Expansion Front of the Parasite Varroa destructor. PLoS Pathog. 2014;10(8).

43. Traynor KS, Rennich K, Forsgren E, Rose R, Pettis J, Kunkel G, et al. Multiyear survey targeting disease incidence in US honey bees. Apidologie. 2016;47(3):325–47.

44. Yuan C, Jiang X, Liu M, Yang S, Deng S, Hou C. An Investigation of Honey Bee Viruses Prevalence in Managed Honey Bees (Apis mellifera and Apis cerana) Undergone Colony Decline. Open Microbiol J. 2021;15(1):58–66.

45. Iqbal J, Mueller U. Virus infection causes specific learning deficits in honeybee foragers. Proc R Soc B Biol Sci. 2007;274(1617):1517–21.

46. Benaets K, Van Geystelen A, Cardoen D, De Smet L, De Graaf DC, Schoofs L, et al. Covert deformed wing virus infections have long-term deleterious effects on honeybee foraging and survival. Proc R Soc B Biol Sci. 2017;284(1848).

47. Coulon M, Dalmon A, Di Prisco G, Prado A, Arban F, Dubois E, et al. Interactions Between Thiamethoxam and Deformed Wing Virus Can Drastically Impair Flight Behavior of Honey Bees. Front Microbiol. 2020;11(April):1–12.

48. Durand AM, Bonjour-Dalmon A, Dubois E. Viral Co-Infections and Antiviral Immunity in Honey Bees. Viruses. 2023;15(5).

49. Sircoulomb F, Dubois E, Schurr F, Lucas P, Meixner M, Bertolotti A, et al. Genotype B of deformed wing virus continues to spread in European honey bee colonies. Submitted to Scientific Reports;

50. Schurr F, Tison A, Militano L, Cheviron N, Sircoulomb F, Rivière MP, et al. Validation of quantitative real-time RT-PCR assays for the detection of six honeybee viruses. J Virol Methods. 2019;270:70–8.

51. Alaux C, Crauser D, Pioz M, Saulnier C, Le Conte Y. Parasitic and immune modulation of flight activity in honey bees tracked with optical counters. J Exp Biol. 2014;217(19):3416–24.

52. Abbott WS. A METHOD OF COMPUTING THE EFFECTIVENESS OF AN INSECTICIDE. J Econ Entomol. 1925;18:265–7.

53. Bailey L. The multiplication and spread of sacbrood virus of bees. Ann Appl Biol. 1969;63(3):483–91.

54. Bailey L, FERNANDO EFW. Effects of sacbrood virus on adult honey-bees. Ann Appl Biol. 1972;72(1):27–35.

55. Locke B, Forsgren E, Miranda JR De. Increased Tolerance and Resistance to Virus Infections: A Possible Factor in the Survival of Varroa destructor-Resistant Honey Bees (Apis mellifera). 2014;9(6).

56. Richard FJ, Aubert A, Grozinger CM. Modulation of social interactions by immune stimulation in honey bee, Apis mellifera, workers. BMC Biol. 2008;6:1–13.

57. Posada-Florez F, Lamas ZS, Hawthorne DJ, Chen Y, Evans JD, Ryabov E V. Pupal cannibalism by worker honey bees contributes to the spread of deformed wing virus. Sci Rep [Internet]. 2021;11(1):8989. Available from: 10.1038/s41598-021-88649-y

58. Moret Y, Schmid-Hempel P. Survival for immunity: The price of immune system activation for bumblebee workers. Science (80-). 2000;290(5494):1166–8.

59. Riessberger-Gallé U, Hernández López J, Schuehly W, Crockett S, Krainer S, Crailsheim K. Immune responses of honeybees and their fitness costs as compared to bumblebees. Apidologie. 2015;46(2):238–49.

60. Bull JC, Ryabov E V., Prince G, Mead A, Zhang C, Baxter LA, et al. A Strong Immune Response in Young Adult Honeybees Masks Their Increased Susceptibility to Infection Compared to Older Bees. PLoS Pathog. 2012;8(12):11–4.

61. Doublet V, Poeschl Y, Gogol-Döring A, Alaux C, Annoscia D, Aurori C, et al. Unity in defence: Honeybee workers exhibit conserved molecular responses to diverse pathogens. BMC Genomics. 2017;18(1):1–17.

62. Amdam G V., Simões ZLP, Hagen A, Norberg K, Schrøder K, Mikkelsen Ø, et al. Hormonal control of the yolk precursor vitellogenin regulates immune function and longevity in honeybees. Exp Gerontol. 2004;39(5):767–73.

63. Amdam G V., Aase ALTO, Seehuus SC, Kim Fondrk M, Norberg K, Hartfelder K. Social reversal of immunosenescence in honey bee workers. Exp Gerontol. 2005;40(12):939–47.

64. Amdam GV, Omholt SW. The hive bee to forager transition in honeybee colonies: The double repressor hypothesis. J Theor Biol. 2003;223(4):451–64.

65. Brutscher LM, Daughenbaugh KF, Flenniken ML. Antiviral defense mechanisms in honey bees. Curr Opin Insect Sci [Internet]. 2015;10(Ccd):71–82. Available from: 10.1016/j.cois.2015.04.016

66. Pizzorno MC, Field K, Kobokovich AL, Martin PL, Gupta RA, Mammone R, et al. Transcriptomic responses of the honey bee brain to infection with deformed wing virus. Viruses. 2021;13(2).

67. Zanni V, Frizzera D, Marroni F, Seffin E, Annoscia D, Nazzi F. Age-related response to mite parasitization and viral infection in the honey bee suggests a trade-off between growth and immunity. PLoS One [Internet]. 2023;18(7 July):1–23. Available from: 10.1371/journal.pone.0288821

68. Corona M, Branchiccela B, Alburaki M, Palmer-Young EC, Madella S, Chen Y, et al. Decoupling the effects of nutrition, age, and behavioral caste on honey bee physiology, immunity, and colony health. Front Physiol. 2023;14(March):1–16.

